# Arabidopsis EID1 E3 ubiquitin ligase regulates acquired thermotolerance by modulating HSBP translocation

**DOI:** 10.1101/2024.12.22.627358

**Authors:** Yi-Tsung Tu, Ting-Chen Yen, Guan-Lin Chuo, Zhi-Qing Wu, Tsung-Luo Jinn, Chin-Mei Lee

**Affiliations:** Institute of Plant Biology, National Taiwan University, Taipei, 106319, Taiwan; Department of Life Science, National Taiwan University, Taipei, 106319, Taiwan; The Master Program in Global Agriculture Technology and Genomic Science, National Taiwan University, Taipei, 106319, Taiwan; International Joint Degree Master’s Program in Agro-biomedicine in Food and Health, National Taiwan University, Taipei, 100233, Taiwan

## Abstract

Climate change is causing a rapid increase in global average temperatures and more frequent heatwaves, posing serious threats to agricultural production and global biodiversity. In response to heat stress (HS), plants can develop acquired thermotolerance (AT) by initiating a heat shock response (HSR) after mild HS priming, thereby enhancing their ability to withstand subsequent later lethal HS events. Central to this process are the HEAT SHOCK FACTORs (HSFs), which form trimeric complexes and activate the expression of *HEAT SHOCK PROTEIN*s (*HSP*s) and other *HSFs* to maintain proper protein and cellular functionality. After heat stress subsides, the HSFs’ activities can be modulated to attenuate the negative effects of HSR during the heat. The SHOCK FACTOR BINDING PROTEIN (HSBP) is a conserved microprotein that plays a prominent role in modulating HSF activities. HSBP can translocate from the cytoplasm into the nucleus during heat stress to directly interact with HSFs and prevent the formation of HSF timers. However, the mechanism that regulates the HSBP cytoplasmic-nuclear shuttling remains unclear. Here, we identified an F-box E3 ubiquitin ligase, EMPFINDLICHER IM DUNKELROTEN LICHT 1 (EID1), whose mutant form shows reduced thermotolerance in AT. We showed that EID1 interacts with HSBP to modulate HSBP cytoplasm-nuclear localization during heat stress, possibly through modulating the K41 of HSBP. The decreased thermotolerance in the *eid1* mutant can be explained by alterations of some *HSPs* expression caused by the mis-localization of HSBP. This finding provided a novel example of E3 ubiquitin-mediated regulation of heat stress in plants.

## Introduction

Due to global warming, the increasing occurrence of high temperatures poses heat stress to plants, which affects plant growth and development and leads to the deterioration of crop quality and yields. In nature, fluctuations in daily temperature shape the plants’ abilities to acclimate to high temperatures. While plants are equipped with innate thermotolerance (or called “basal thermotolerance”, BT) to cope with heat stress, some plants can boost their ability through a high but not lethal temperature, called heat priming (Yeh et al., 2012). After priming, plants acquire heat memory to sustain their ability to fight further lethal high temperatures. Depending on the time that the memory can last, the acquired thermotolerance (AT) can be classified into short-term acquired thermotolerance (SAT) and long-term acquired thermotolerance (LAT), depending on whether the memory can last shorter or longer than 48 hours after heat priming. The other class is long-term moderate high-temperature responses (TMHT), which measures the plants’ thermotolerance during prolonged moderate high temperatures.

The heat stress regulatory network is complicated, and different key regulators are needed to defeat heat stress (Balazadeh, 2022; Saini et al., 2022). Upon heat stress, a group of HEAT SHOCK FACTORs (HSFs) can be induced and subsequently regulate downstream *HEAT SHOCK PROTEINS* (*HSPs*) and other heat stress response (HSR) genes to help plants survive under heat stress. Among the 21 Arabidopsis HSFs, the Class A HSFs positively regulate HSRs, while Class B HSFs function as negative regulators of HSRs. The functions of Class C HSFs are not fully understood. The most well-characterized regulator is HSFA1, a major early heat response regulator that activates the expression of multiple HSRs. In this process, the inactive HSFA1 monomers translocate from the cytoplasm to the nucleus and form active HSF trimers to interact with the promoters of HSRs. These HSR genes encode HEAT SHOCK PROTEIN (HSPs), LEA proteins, increasing amino acid contents, anti-oxidants, and chaperones to alleviate the oxidative and protein misfolding toxicities to the cells (Saini et al., 2022). Other HSFs can function as positive or negative regulators during heat stress. In addition to HSFs, DEHYDRATION-RESPONSIVE ELEMENT BINDING PROTEIN 2A (DREB2A) transcription factor induced by HSFA1s acts as a positive regulator of HSR (Yoshida et al., 2011). Another transcription factor induced by HSFA1s, MULTIPROTEIN BRIDGING FACTOR 1C (MBF1C), activates both the HSR positive regulator, *DREB2A*, and negative regulator, *HSFBs* (Suzuki et al., 2011; Yoshida et al., 2011). In turn, HSFBs feedback suppresses the transcription of HSFA1s, as part of the complicated regulatory network in heat stress (Ikeda et al., 2011).

When the heat stress ceases, or during the recovery stage after heat priming in AT, the activity of HSFA1 can be attenuated by HSPs and other negative regulators. The microprotein HEAT SHOCK FACTOR-BINDING PROTEIN (HSBP) has been reported as one of the negative regulators by interacting with HSFA1 to inactivate it by interacting with them to hinder the formation of HSFA1 trimers during SAT (Hsu et al., 2010). HSBP is conserved in many species, including in animals and plants (Tai et al., 2002). In humans, HSBP has been shown to be localized in the nucleus and interact with Hsf1, negatively regulating the HSR by inhibiting DNA-binding and transactivation activities of Hsf1 (Satyal et al., 1998). Different from human, Arabidopsis HSBPs originally localize in the cytoplasm under normal temperature, translocate into nucleus upon heat stress, and then relocate back to cytoplasm during recovery (Hsu et al., 2010; Huang et al., 2024). The proper translocations of HSBP into the nucleus are critical for inactivating HSFA1s and HSFA2, and the translocation of and interaction functions both lies in the coiled-coil domains of HSBP (Hsu et al., 2010; Hsu and Jinn, 2010). Although *hsbp* mutant only increased thermotolerance in SAT but not BT, recent reports demonstrate that HSBP also interact with other class A, class B and class C HSFs (Fu et al., 2006; Hsu et al., 2010; Huang et al., 2024). These HSFs can act as both positive and negative heat stress regulators, and HSBP can regulate transcript levels of some *HSFs*, indicating more complicated regulatory mechanism of HSBP in heat stress. While the negative regulatory roles of HSBP in heat stress seem conserved across the kingdoms, the mechanism for its translocation is still understudied.

Protein ubiquitination plays a role in modulating heat stress regulator and their functions (Lim et al., 2013; Liu et al., 2016; Peng et al., 2019; Kim et al., 2020; Zhang et al., 2021). In this process, E3 ubiquitin ligases interact with their specific substrates and transfer ubiquitin onto their substrates (Hua and Vierstra, 2011). Several E3 ubiquitin ligases have been reported to affect thermotolerance. Overexpression of *Arabidopsis PUB48*, rice *HCI1,* rice *HTAs,* or *Solanum CHIP* increases thermotolerance, while overexpression of OsDHSRP1 causes hypersensitive to heat stress (Lim et al., 2013; Liu et al., 2016; Peng et al., 2019; Kim et al., 2020; Zhang et al., 2021). However, their ubiquitinated substrates were not identified. The best characterized E3 ubiquitin ligase and substrate pair are the BTB/POZ and math domain proteins (BPMs), which interact with DREB2A for its ubiquitination and degradation (Morimoto et al., 2017). Additionally, it has been reported that ploy-ubiquitin has largely accumulated under heat stress, but their potential roles and mechanisms remain to be explored (Sharma et al., 2021).

Previously, Luhua et al. (2013) reported that an F-box E3 ubiquitin ligase EMPFINDLICHER IM DUNKELROTEN LICHT 1 (EID1) can be induced by heat and enhances root elongation under TMHT conditions in the *eid1* mutant (called *eid1-1* in this manuscript). EID1 contains an F-box domain in its N-terminus, which has been shown to interact with Skp1-like proteins (ASKs) to form SCF complex (Marrocco et al., 2006). It also contains a leucine-zipper domain (L-ZIP), PEST domain, WSL motif, and EID-like protein domain (ELP) for potential functions in protein-protein interactions and stabilities. It has been demonstrated that EID1 is involved in the phyA-mediated light signaling pathway (Büche et al., 2000; Dieterle et al., 2001; Zhou et al., 2002; Marrocco et al., 2006). Additionally, Müller et al. further showed that the Solanum EID1 decelerates the circadian clock through PHYB1-mediated light input pathways by genetic analyses (Müller et al., 2018). In a recent report, soybean EID1 interacts with evening complex proteins, clock regulators, to stabilize ELF3 protein to affect flowering time and yields (Qin et al., 2023). These results indicate that EID1 may modulate light signaling pathways to regulate the plant circadian clock. While the reports started to unveil the function of EID1 in photoperiod and the circadian clock, their mechanism in regulating thermotolerance remains to be investigated (Luhua et al., 2013).

Here, we demonstrated that the *eid1* mutants exhibited a decrease in SAT measured by heat-induced hypocotyl elongation and survival rate. We further showed that EID1 interacts with the SAT negative regulator, HSBP. Furthermore, EID1 regulates HSBP translocations from the cytoplasm to the nucleus, potentially through ubiquitination of the lysine 41 of the HSBP, which may partially explain the thermotolerance phenotypes in the *eid1* mutants. These findings uncover the mechanism by which EID1 regulates thermotolerance and the role of E3 ubiquitin ligases in heat stress.

## Results

### The *eid1* mutants are compromised for the basal (BT) and short-term acquired thermotolerance (SAT)

To investigate the mechanism that EID1 regulates thermotolerance, we acquired three *eid1* mutant alleles, including previously identified *eid1-1* (SALK_021058), and two other alleles, *eid1-2* (SALK_013061) and *eid1-3* (SALK_027403) (**Supplemental Figure S1B-S1C**). Among them, the previously reported *eid1-1* is a knockdown mutant, producing residue transcripts after the T-DNA insertion site. Therefore, we used *eid1-2* and *eid1-3* knockout mutant alleles to further characterize their thermotolerance phenotypes (**Supplemental Figure S1B-S1D**).

We first examined the thermotolerance phenotypes of *eid1* mutants (**Figure 1**). Both *eid1-2* and *eid1-3* exhibited decreased heat-induced root elongation compared to wildtype under basal thermotolerance (BT) and the short-term acquired thermotolerance (SAT) conditions (**Figure 1A-1B**). We further examined the survival rates under BT and SAT, *eid1-2* and *eid1-3* also showed decreased overall survival rate compared to Col-0 (**Figure 1C-1D**). To further examine whether EID1 affect the expression of HSR genes, we analyzed the expression of some HSFs and HSPs during the heat treatment and recovery regimes (**Figure 2**). Although the heat-induced expression of *HSP101*, *HSP70*, *HSP17.6A* and *HSP18.2* were overall decreased in the *eid1* mutant compared to Col-0 after 1 hour in 37°C (HS) and after recovery 1 hour at 22°C (R1); however, the differences were not significant. On the other hand, the expression of *HSFA7a*, *APX2*, and *HSP17.6II* were significantly decreased in the *eid1* mutant under HS and R1 conditions. Among them, the expression of *APX2* were still decrease after 2 hours during recovery (R2). Additionally, another small HSP, *HSP17.6C* exhibited significant decreased in the *eid1* mutant under HS. These results indicate that the EID1 function as a positive regulator in the BT and SAT responses.

**Figure 1.**
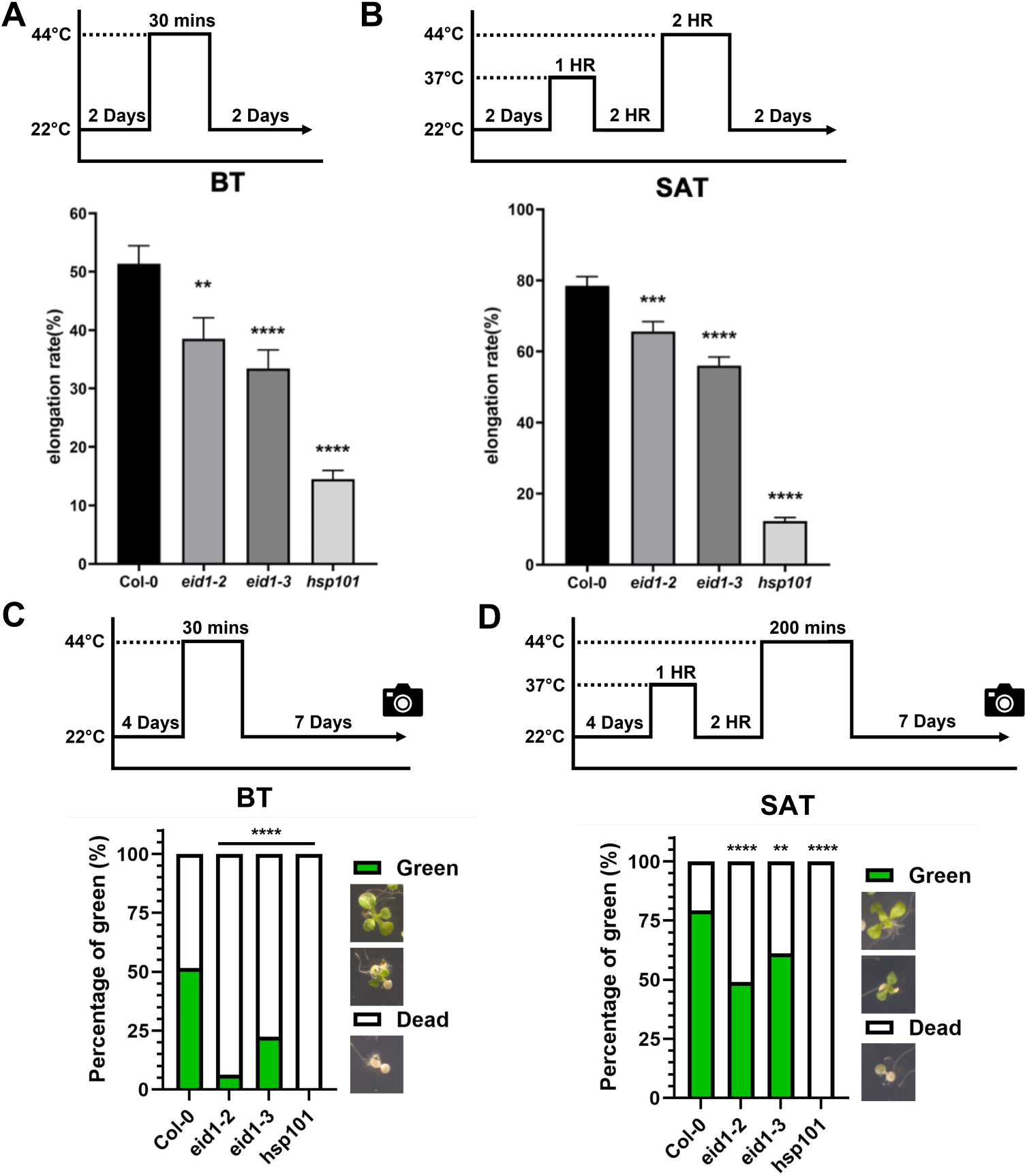
The *eid1* mutants exhibit the heat-treated phenotype in hypocotyl elongation and survival rate. **(A)** The 2-d-old seedlings of Col-0 and *eid1* mutants were treated at 44°C for 30 min, following 2 days of recovery (BT treatment) as schematically shown on the top panel. The hypocotyl elongation rates were measured. (\*\**p*<0.01, \*\*\*\**p*<0.001, two-tailed *t*-test) **(B)** The 2-d-old seedlings of Col-0 and *eid1* mutants were treated with SAT treatment and recovered for 2 days, as schematically shown on the top panel, and then the hypocotyl elongation rates were measured. **(C)** The survival rate of *eid1* mutants in BT treatment. The 4-d-old seedlings of *eid1* mutants were treated at 44°C for 30 min following 7 days of recovery (BT treatment), as schematically shown on the top panel. The survival rates were calculated after recovery. **(D)** The survival rate of *eid1* mutants in SAT treatment. The 4-d-old seedlings of *eid1* mutants were treated at 37°C for 1 hr and recovered for 2 hr, challenged at 44°C for 200 min, and then recovered for 8 days before the survival rates were calculated, as the scheme shown on the top panel. The data are means of three biological repeats. Error bars represent SD. (chi-square test with Bonferroni correction, ** *p*<0.01, **** *p*<0.001

**Figure 2.**
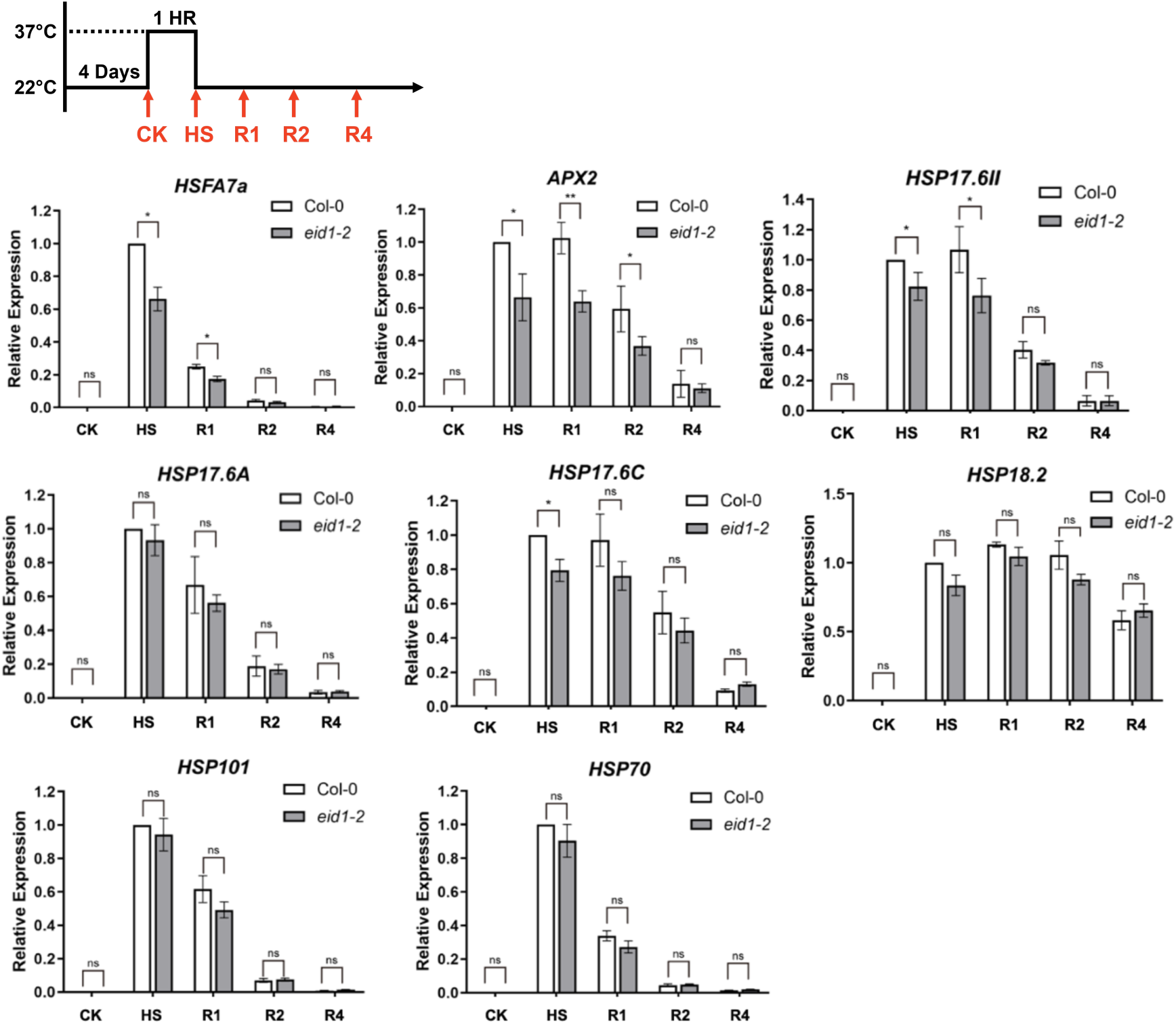
The expression of heat-responsive genes in Col-0 and *eid1* mutant during heat stress. The 4-d-old seedlings of Col-0 and *eid1-2* mutant were treated with SAT condition (scheme on the top panel) and analyzed by real-time qRT-PCR. (ns, not significant, \**p*<0.05, \*\**p*<0.01, two-tailed *t*-test)

### The subcellular localizations of EID1 can be regulated by high temperature

We further determined whether the expression of *EID1* can be regulated by high temperature and recovery (**Figure 3**). The mRNA expression of *EID1* during the SAT was not significantly induced by 37℃ HS or recovery for 1, 2, or 4 hours (R1, R2, and R4) (**Figure 3A**). The GUS reporters driven by the *EID1* native promoter, *EID1* mainly expressed in the vascular tissue in the Arabidopsis seedlings under normal temperature (**Figure 3B**). Upon heat priming, the expression of *EID1* slightly spread through the leaf tissues. While the recovery stage, the GUS signals were decreased in the leaf and vascular tissues. We also determined whether the EID1 protein is regulated post-translationally (**Figure 3C**). The Arabidopsis seedlings expressing *EID1-GFP* driven by its native promoter were treated with HS and then recovery, and the EID1-GFP protein levels did not significantly change under the heat regimes tested. We further examined the subcellular localization of the EID1 protein (**Figure 3D**). The EID1 protein localized in both cytoplasm and nuclei under normal temperature, and under HS and recovery, the EID1 proteins mainly localized in the nuclei. These results suggested that EID1 is not regulated by heat transcriptionally or translationally, but the protein localizations can be modulated by heat.

**Figure 3.**
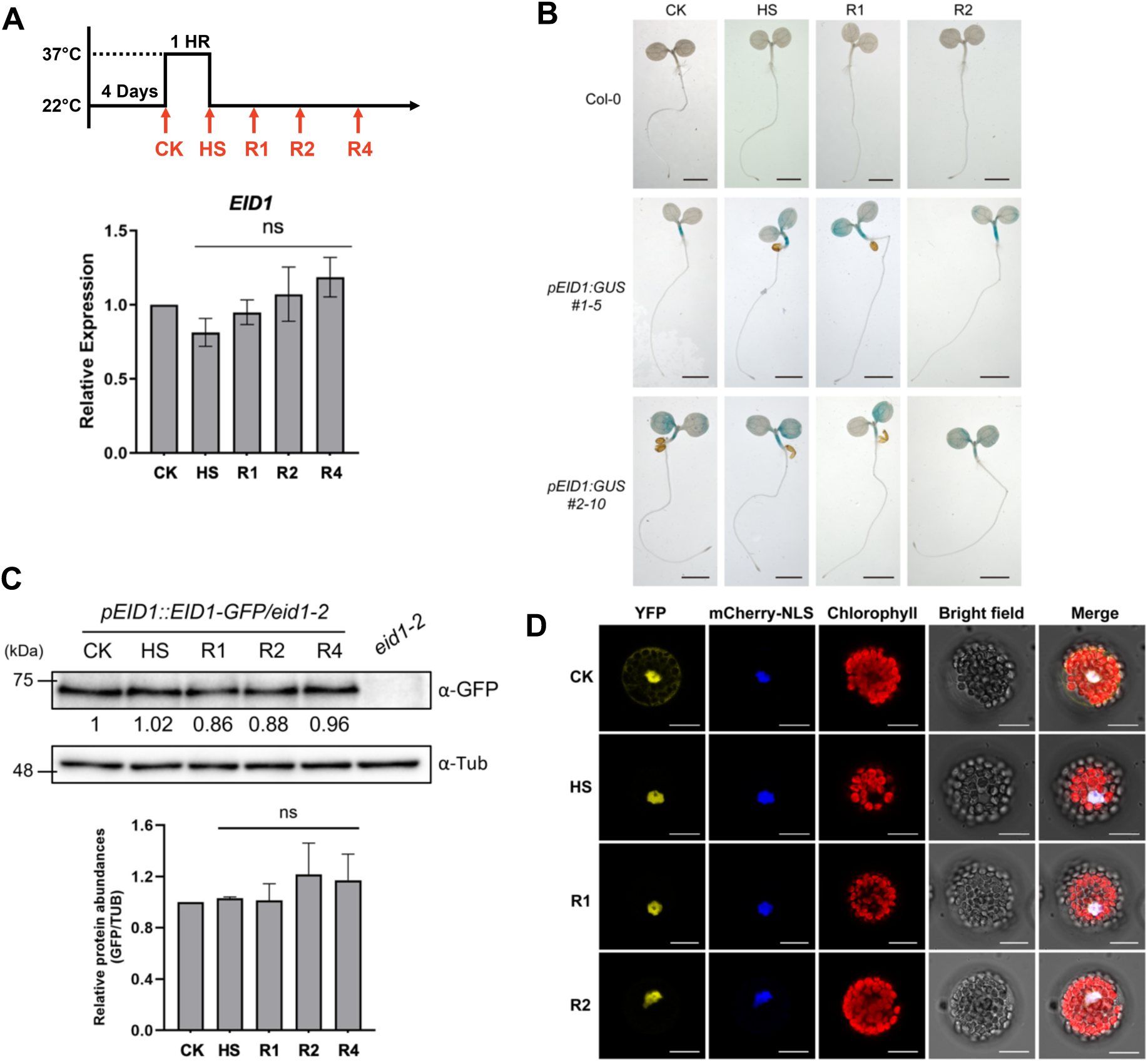
The transcripts and protein level of EID1 after heat treatment. **(A)** The transcript level of *EID1* at SAT condition (scheme on the top panel) in 4-d-old seedlings of Col-0 as measured by real-time qRT-PCR. (ns, not significant, two-tailed *t*-test) **(B)** GUS staining of *pEID1::GUS* under heat stress and recovery stage. 4-day-old *pEID1::GUS* seedlings were stressed at 37 °C for 1 hr (HS) and then recovered for 1 or 2 hours (R1 or R2). **(C)**The protein levels of EID1 under heat stress and recovery stages were assessed by immunoblot analysis. The 4-d-old *pEID1::EID1-GFP/eid1* were stressed at 37 °C for 1hr (HS), and following recovered 1, 2, or 4 hours (R1, R2, and R4). EID1-GFP was recognized by an anti-GFP antibody, and tubulin was used as a loading control. The protein levels were quantified using ImageJ software, and three biological repeats were shown on the lower panel. (ns, not significant, two-tailed *t*-test) **(D)** EID1 cellular localization under heat stress. YFP-EID1 and mCherry-NLS were cotransfected in Col-0 protoplasts and treated at 37°C for 1 hour (HS), and following recovered for 1 or 2 hours (R1 or R2). The fluorescence signals were observed by a confocal microscope. Bar= 20μm. The data were means of three biological repeats. Error bars represent SD.

### EID1 regulates SAT through SAT negative regulator, HSBP

We intended to explore the mechanism by which EID1 regulates SAT by identifying the substrates of EID1. In considering the positive regulatory role of EID1 in SAT (**Figure 1**) and the E3 ubiquitin ligase function of the EID1 protein. We hypothesized that EID1 might target negative regulators in the thermotolerance pathways. Therefore, a negative regulator of HSFs, the HSBP, whose mutation can cause an increase in the SAT, was chosen. To test this hypothesis, we performed BiFC and co-immunoprecipitation (co-IP) assays and found that EID1 interacts with HSBP in plants (**Figure 4A-4B**). To further determine whether HSBP is responsible for the decrease of SAT in the *eid1* mutant, we generated the *eid1/hsbp* double mutant and compared its survival rate in SAT to single mutants (**Figure 4C**). The *hsbp* mutant slightly increased its survival rate while the *eid1* mutant decreased it. The double mutant exhibited similar pattern to *hsbp* mutant, indicating *HSBP* epistatic to *EID1* in SAT.

**Figure 4.**
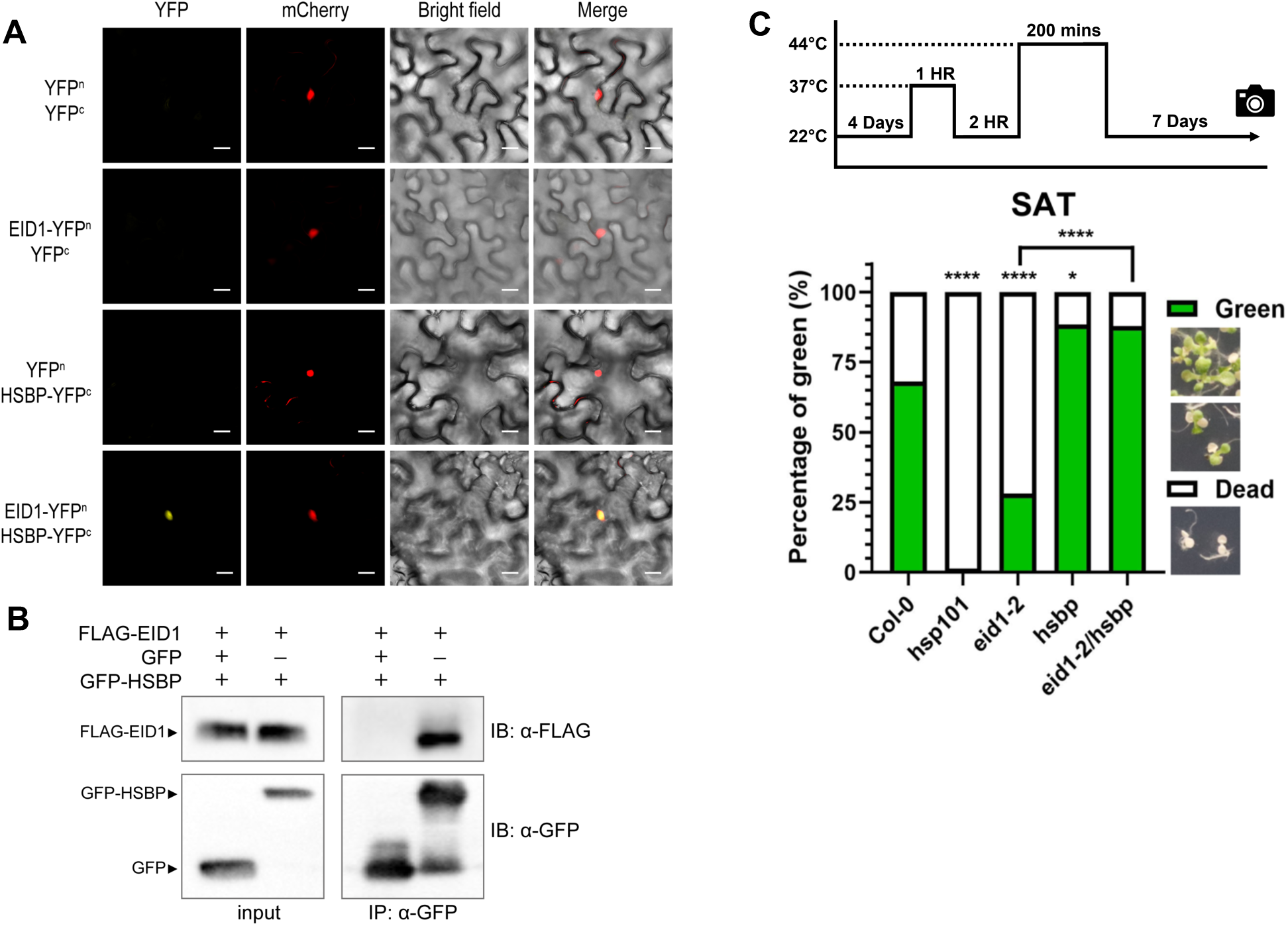
The interaction between EID1 and HSBP by BiFC assay and co-IP assay. **(A)** BiFC assay of EID1 and HSBP in tobacco leaves. HSBP and EID1 were fused to the N-terminal (nYFP) and C-terminal (cYFP) of YFP, respectively. HSBP-nYFP, EID1-cYFP, and mCherry-NLS were coexpressed in tobacco leaves. The mCherry-NLS was used as a nuclear marker. Bar= 20μm. **(B)** Co-IP assays of EID1 and HSBP in tobacco leaves. HSBP and EID1 were fused to GFP and FLAG, respectively. HSBP-GFP and FLAG-EID1 were coexpressed in tobacco leaves. The anti-GFP antibody was used to immunoprecipitate HSBP-GFP. The anti-GFP and anti-FLAG antibodies were used for immunoblot analysis. **(C)** The HSBP is epistatic to EID1 in SAT. The survival rate of *eid1*, *hsbp*, and *eid1/hsbp* mutant in SAT treatment. The 4-day-old seedlings were treated at 37°C for 1 hour and recovered for 2 hours, challenged at 44°C for 200 min, then recovered for 7 days before the survival rates were calculated. The scheme was shown on the top panel. (chi-square test with Bonferroni correction, * *p*<0.05, **** *p*<0.001). The data are means of three biological repeats. Error bars represent SD.

### EID1 regulates cytoplasm-nuclear localization of HSBP

To investigate the molecular mechanism of how EID1 regulates SAT through HSBP, we first measured the HSBP transcript levels in the *eid1* mutants, and the HSBP transcript levels were similar to that in the wild type (**Supplemental Figure S2**).

Previously, Hsu et al. (2010) reported that HSBP can shuttle from the cytoplasm into the nucleus to interact with HSFs and attenuate HSF activities during the heat priming and recovery phase. We, therefore, determined the HSBP-GFP protein localization in the Col-0 and *eid1* mutants. The HSBP-GFP increased in the nucleus after 1 hour of 37℃ treatments (HS) and then gradually decreased throughout 1 and 2 hours of recovery (**Figure 5, top panel**). In the *eid1* mutants, the signals of HSBP-GFP were less in the nucleus under HS or recovery stages (**Figure 5, lower panel**). These data indicated that decreased translocation of HSBP in the *eid1* mutant may lead to the misregulation of HSF activities and a decreased SAT.

**Figure 5.**
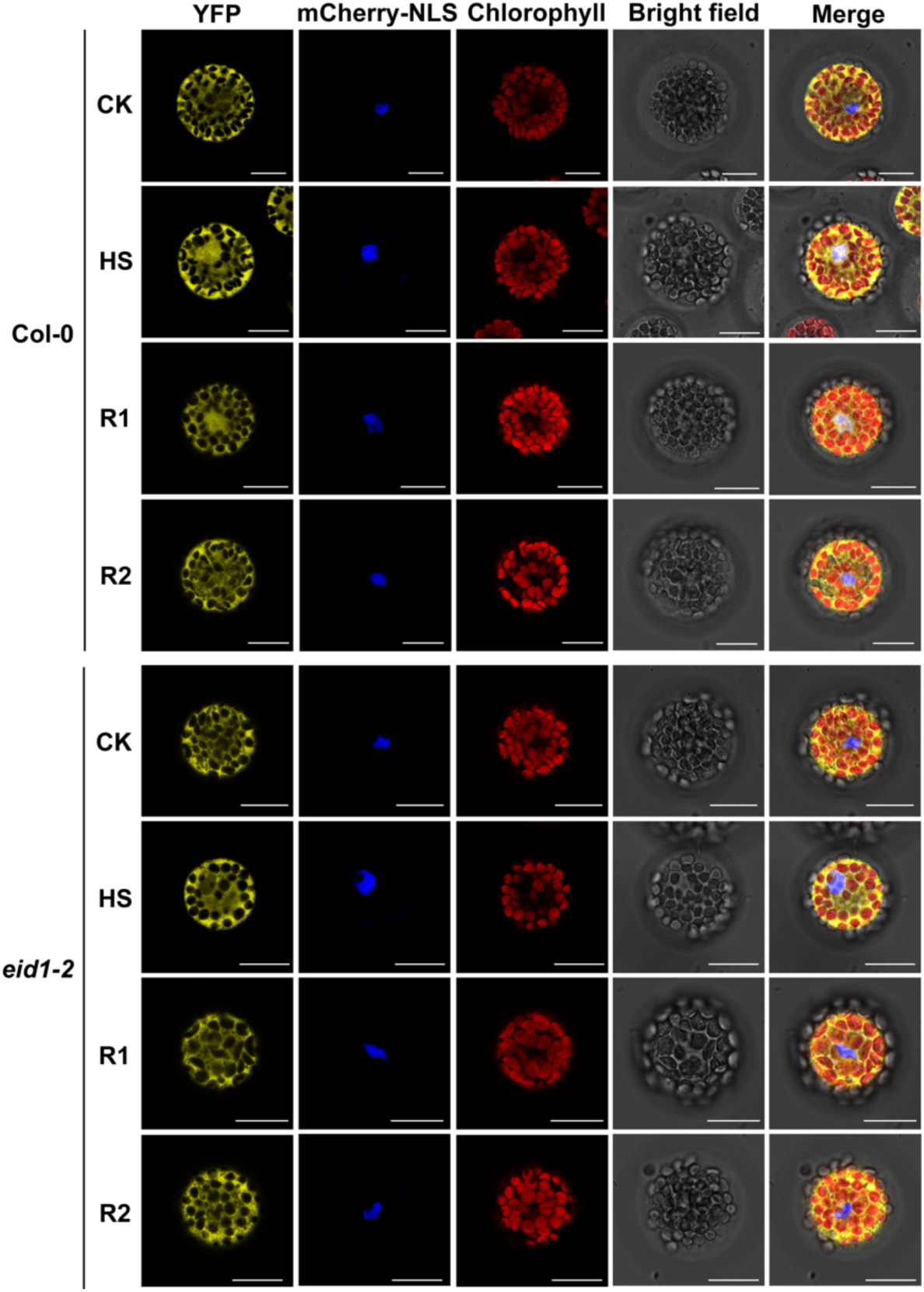
EID1 affects the translocation of HSBP to the nucleus in response to heat. Translocation of the HSBP protein in the Col-0 and *eid1-2* protoplasts under heat stress. YFP-HSBP and mCherry-NLS were cotransfected in Col-0 and *eid1-2* protoplasts and treated at 37°C for 1 hour (HS), and following recovered 1 or 2 hours (R1 or R2). The fluorescence signals were observed by a confocal microscope. Bar= 20μm.

We further examine whether the potential ubiquitination lysine (K) sites of HSBP can affect their subcellular localization, so we mutated the K10, K41, K76, and K80 to Arginine (R) and observed their translocation to the nucleus during heat treatment (**Figure 6**). While HSBP-K74R-GFP and HSBP-K80R-GFP exhibited similar cytoplasm-nucleus translocations patterns to wildtype during heat stress and recovery. The HSBP-K41R-GFP showed fewer signals in the nucleus under heat stress and recovery, and the nuclear-localized HSBP-K10R-GFP in the recovery 1 hour (R1) largely decreased compared to that in the wildtype. Whether the EID1 affects HSBP cytoplasm-nuclear shuttling through ubiquitination of the K41, and maybe K10 as well, in HSBP remains to be validated.

**Figure 6.**
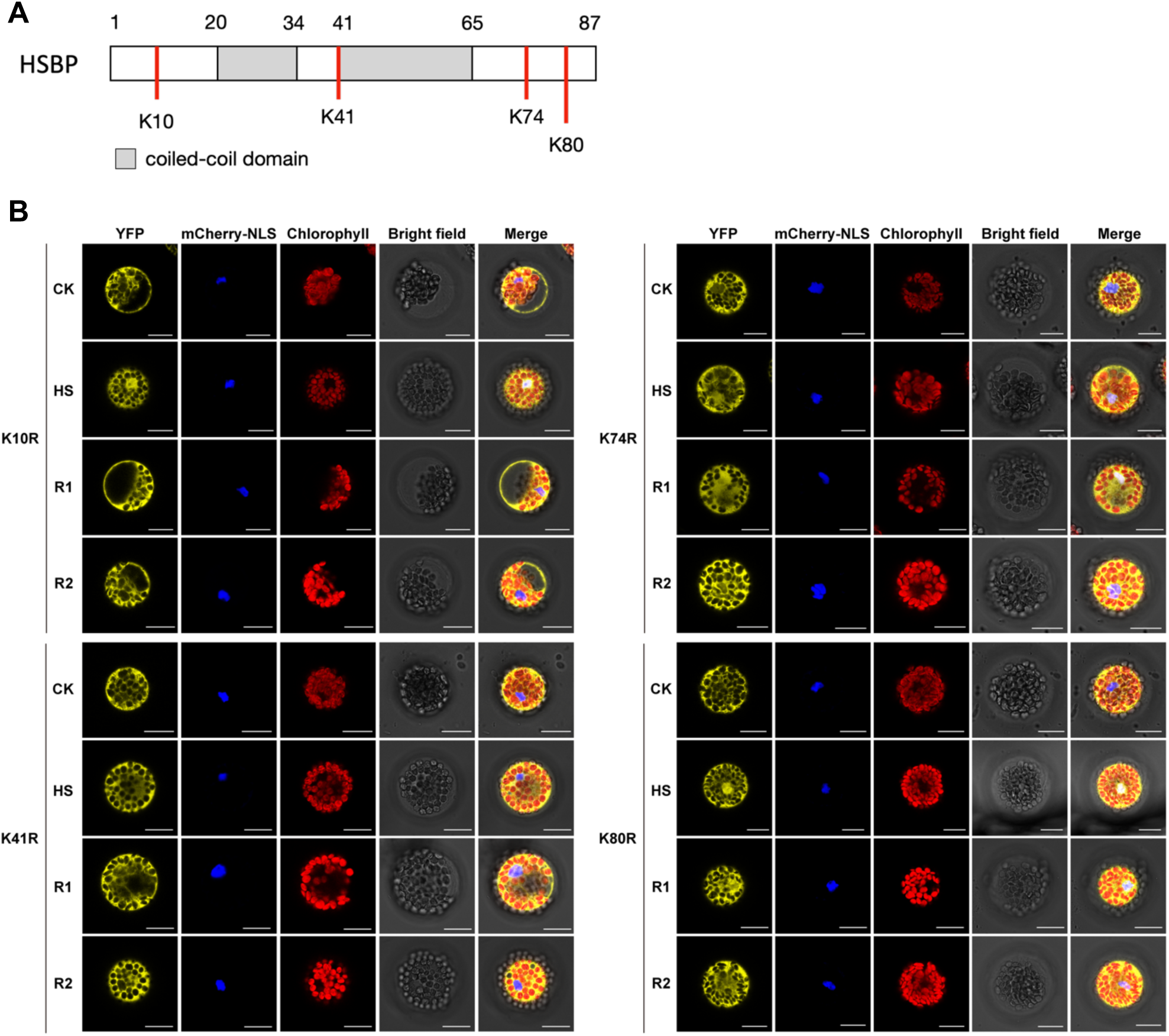
The K41 site mutation affects the translocation of HSBP under heat stress. **(A)** The diagram showing Lys positions in the EID1 protein. The number, gray boxes, and red lines represent amino acid, coiled-coil domain, and Lys positions, respectively. **(B)** The subcellular localization of HSBP with K mutation under heat stress. The Lys of HSBP was mutated to Arg and fused with YFP tag. The HSBP mutation and mCherry-NLS were co-transfected in Col-0 protoplasts and treated at 37°C for 1 hour (HS), and following recovered 1 or 2 hours (R1 and R2). The fluorescence signals were observed by a confocal microscope. Bar= 20 μm.

## Discussion

Protein ubiquitination is a ubiquitous post-translational regulatory mechanism in regulating board biological pathways, including heat stress. Although several E3 ubiquitin ligases have been reported to regulate heat stress. Not until recently have BPMs been demonstrated to regulate the protein stabilities of DREB2A (Morimoto et al., 2017). This study uncovered the thermotolerance functions and mechanism of the previously identified heat stress regulator, EID1 (Luhua et al., 2013). We showed that loss of *EID1* can cause a decrease in short-term acquired thermotolerance (SAT) and basal thermotolerance (BT) (**Figure 1**). These phenotypes can be partially explained by the alterations of some HSR genes (**Figure 2**). To determine the mechanism by which EID1 regulates SAT, we found that EID1 can interact with HSBP, an SAT-negative regulator, which can translocate into the nucleus to deactivate HSF functions (**Figure 5**). Additionally, a larger portion of HSBP protein is distributed in the cytoplasm than in the nucleus in the *eid1* mutant, indicating EID1 regulates HSBP translocation from the cytoplasm to the nucleus. This translocation can be totally eliminated by a mutation in the lysine 41, a potential ubiquitination site of HSBP (**Figure 6**). These results revealed that EID1 regulates HSBP translocations during SAT through protein ubiquitination, and it in turn to affect the complex HSF-mediated heat stress regulatory networks (**Figure 7**).

**Figure 7.**
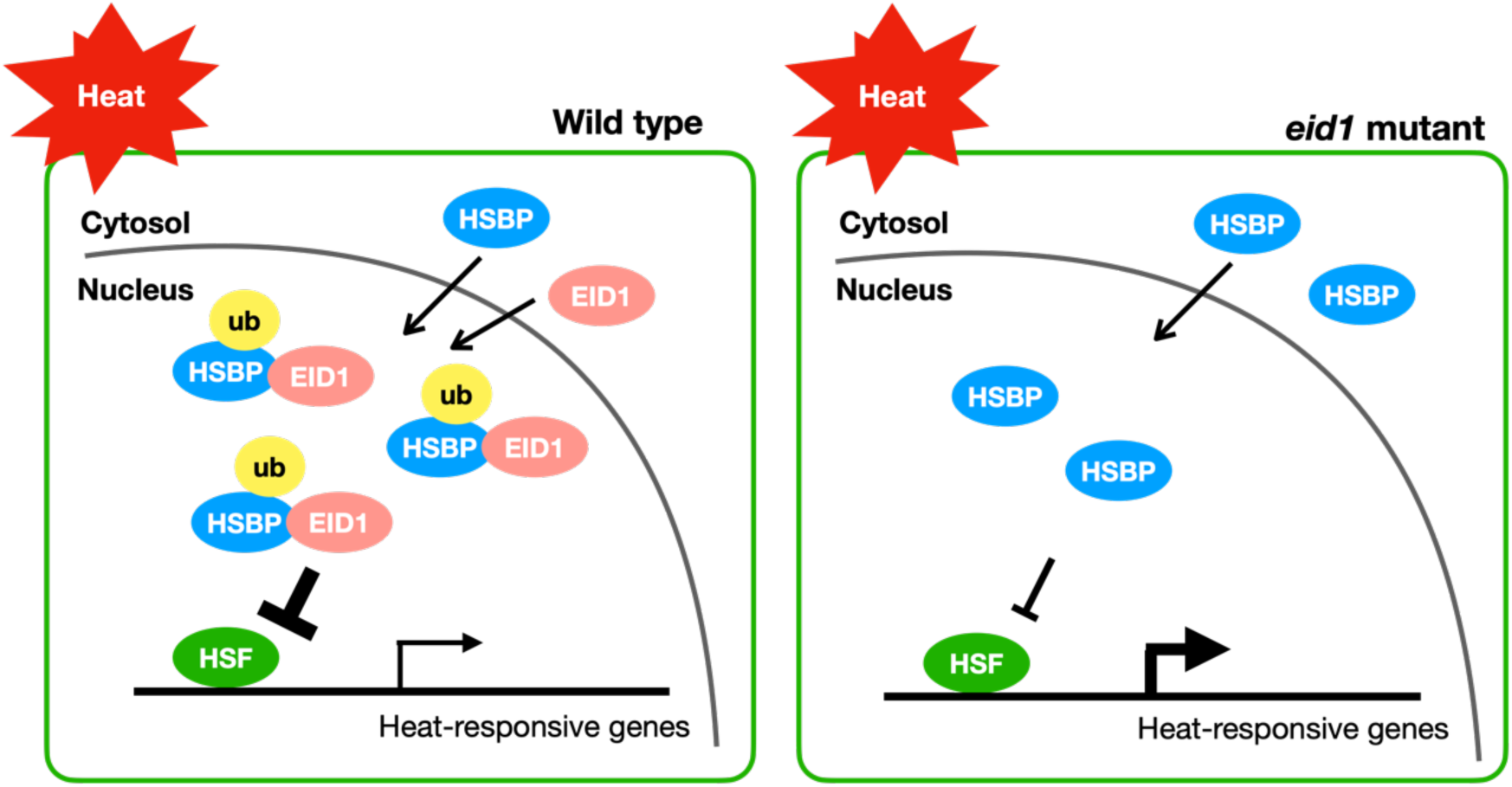
The working model of EID1 regulates heat response by modulating HSBP translocation. EID1 interacts with HSBP to mediate HSBP protein expression and localization, possibly through a putative ubiquitination site on HSBP, thereby regulating HSF gene expression in response to heat stress. In the *eid1* mutant, HSBP translocation is inefficient in regulating the heat stress response.

### EID1 regulates HSBP cytoplasm-nucleus shuttling

EID1, as a *bona fide* E3 ubiquitin ligase shown to form SCF complex, has the potential to ubiquitinate HSBP (Marrocco et al., 2006). Proteins with ubiquitin modifications can lead to several consequences: protein degradation, changes in protein activities, or subcellular localization. The EID1 localized in both cytoplasm and nuclei under normal temperature and translocated into nuclei under heat stress and recovery (**Figure 3D**). The EID1 also can regulate the subcellular localization of HSBP proteins (**Figure 5-6**). The HSBP protein distribution in the nucleus is less than that in the wild type. Additionally, the retention of the HSBP proteins during the recovery phase dropped faster in the *eid1* mutant, suggesting the EID1 is required to regulate the HSBP shuttling between the cytoplasm and nucleus. Furthermore, the mutation of the putative ubiquitination site, lysine (K) 41 of the HSBP, affects its translocation to the nucleus during heat stress (**Figure 6**). While mutation in the K10 of HSBP increases its relocation back to cytoplasm in the recovery stage. The K63 mono-ubiquitination has been reported as the protein cargo-sorting ubiquitin mark (Romero-Barrios and Vert, 2018). Whether K41 and K10 of HSBP can be conjugated with K63-linked ubiquitin by EID1 will be worthy of examination. Furthermore, the thermotolerance of SAT of K41-mutated HSBP can be investigated. The K41 located in the coiled-coil domain of HSBP (**Figure 6A**), which is critical for HSBP translocating into nuclei and interacting with HSFs as previous reports (Hsu et al., 2010; Hsu and Jinn, 2010; Huang et al., 2024). The ubiquitination of K41 of HSBP may affect the coiled-coil structure and/or the interaction between HSBP and HSFs, affecting thermotolerance.

### EID1 can play positive or negative roles in heat stress

The *eid1* mutants exhibited decreased thermotolerance in SAT and BT (**Figure 1**) and enhanced thermotolerance under TMHT (Luhua et al., 2013), implying EID1 may be involved in different heat regimes or stages of heat stress. In our study, we found EID1 interacted with HSBP and affected its protein translocation into the nucleus during heat stress. The proper cytoplasmic and nuclear shuttling of HSBP during heat stress is critical for HSBP to interact and repress HSFs’ activities (Hsu et al., 2010). Increasing evidences showed that HSBP interacts and regulates more than heat stress positive regulators, HSFA1s (Fu et al., 2006; Hsu et al., 2010; Huang et al., 2024). Recently, our findings showed that HSBP interacts with 16 HSFs, including the A class HSFs as HSR positive regulators and class B HSFs for HSR negative regulators (Huang et al., 2024). These Class A, Class B and Class C HSFs, which can pose both positive or negative effects in the establishments of thermotolerance. As EID1 affects the translocations between cytoplasm and nuclei (**Figure 5 and 6**), which makes the overall effects on EID1 more complicated. The translocations of HSBP tightly link to its regulation upon its interacting HSFs and their downstream genes form a complicated regulatory network (Huang et al., 2024). It may also be corroborated by the controversial thermotolerance phenotypes observed in previous and our studies. This may explain not all the HSR genes were affected in the *eid1* mutants in the SAT condition we tested (**Figure 2**). Though the complication, we found *HSFA7a*, *APX2*, *HSP17.6II*, and *HSP17.6C* expression were significantly decreased in the *eid1* mutant under HS and recovery, which may contribute to the decrease of thermotolerance in the *eid1* mutant. Further dissect the heat stress pathways regulated by EID1 through HSBP and its other potential interactors in heat stress remains to be investigated.

## Materials and Methods

### Plant Materials and Growth Conditions

All *Arabidopsis thaliana* lines utilized in this study were derived from the Columbia-0 (Col-0) genetic background. The single mutants *eid1-2* (SALK_013061), and *eid1-3* (SALK_027403) were obtained from the Arabidopsis Biological Resource Center (ABRC). The *hsbp-1* (SALK_081104) (Hsu et al., 2010), and *hsp101* (SALK_066374) (Charng et al., 2006) were the gift from Dr. Tsung-Luo Jinn and Dr. Yee-Yung Charng, respectively. The double mutants *eid1/hsbp* were generated by crossing *eid1-2* and *hsbp-1*.

Arabidopsis plants were grown in the growth chamber at 22°C under long-day conditions (16 hours light/ 8 hours dark) and at a light intensity of 85∼120 μmol m^-2^ s^−1^.For thermotolerance assay, the seeds sterilized with 70% ethanol and 0.1% TRITON® X-100 were stratified at 4°C for 3 days and sowed on the 1/2 strength Murashige and Skoog (MS) medium (Caisson, MSP02) supplemented with 0.8% agar (Duchefa Biochemie, P1003) and 1 % sucrose. For hypocotyl elongation, seedlings were grown on 1/2 MS medium with 1% agar in the dark at 22°C for 2 days. For seed stratification, all seeds were kept at 4°C in the dark for 3 days and then grown on soil.

### Thermotolerance assays and heat treatments

Four-day-old seedlings were grown on 1/2 MS medium before heat treatment. The plates were sealed with plastic electric tape and soaked in the water bath at the indicated temperature (37°C or 44°C) and duration for basal thermotolerance assay (BT) and short-term thermotolerance assay (SAT). The plates were placed back in a 22°C growth chamber during the recovery period before exposure to lethal heat shock at 44°C. The survival rate was determined after a 7-day recovery period. The seedlings with all green leaves and newly developing leaves were counted as “green”, and those with whitening leaves were indicated “dead”.

For hypocotyl elongation assays, the hypocotyl length of a subset of 2-day-old seedlings was measured as “untreated control”, and the remaining subset was incubated in the water bath for BT or SAT as indicated temperature and duration. After 2-day recovery at 22°C in growth chambers, the hypocotyl elongation rate was examined by the equation: [(each treated hypocotyl length (4d)- mean of untreated control)/mean of untreated control].

### Reverse Transcription Quantitative PCR (RT-qPCR)

For RNA extraction, Arabidopsis thaliana seedlings were harvested, flash-frozen in liquid nitrogen, and ground into a fine powder using freezer mill (Retsch, MM400). Total RNA was then extracted from the tissue powder using TRIzol Reagent (Invitrogen, 15596018) following the manufacturer’s instructions and treated with DNase (Promega, M6101). For RT, cDNA was synthesized using PrimeScript RT Reagent Kit (Takara, RR014B). RT-qPCR was performed on CFX Connect Real-Time System (Bio-Rad) with iQ SYBR Green Supermix (Bio-Rad, 170-8880). Expression levels were normalized to the internal control gene *PP2AA3* (AT1G13320). Three independent biological replicates were conducted for each experiment. The sequences of all primers used for RT-qPCR are provided in Supplemental Table S1.

### Protein Extraction and Immunoblot

Protein was extracted from Arabidopsis thaliana seedlings, previously ground to a powder using a freezer mill (Retsch, MM400) in liquid nitrogen. The extraction buffer contained 4 M Urea, 100 mM Tris-HCl (pH 6.8), 5% SDS, 15% Glycerol, supplemented with 0.4% protease inhibitor cocktail (Roche, 04693132001), 0.5% β-mercaptoethanol (PanReac AppliChem, A11080100), and 0.2% bromophenol blue dye (Sigma, 62625289). The extracted protein was separated by sodium dodecyl sulfate-polyacrylamide gels (SDS-PAGEs) and subsequently transferred to Polyvinylidene difluoride (PVDF) membranes (Millipore, Cat. no. IPVH00010) for immunoblotting.

The membranes were probed with a 1:4000 dilution of anti-tubulin antibody (Sigma, T5168) or 1:1000 dilution of anti-GFP (Santa cruz, sc-9996) in the blocking solution for overnight incubation at 4°C, followed by incubated with mouse IgG-conjugated with HRP (1:5000, Promega, W4021) at room temperatures for 1 hr. Signal detection was done using Clarity Western ECL Substrate (Bio-Rad, Cat. no. 1705061) with the iBright™ CL750 Imaging System (Invitrogen). The protein band intensity was quantified using ImageJ software (Schneider et al., 2012). This procedure was independently replicated with three biological samples.

### Bimolecular fluorescence complementation assay

The coding region of *HSBP* and *EID1* were subcloned into *pEarleyGate201-YN* or *pEarleyGate202-YC* (Lu, et al., 2010), respectively, to generate YFP N-terminus or C-terminus fusions constructs and then transformed into the Agrobacterium GV3101 strain. The *pEG202-NLS-mCherry* was co-expressed as a nuclear marker, and the empty vectors were used as negative controls. The overnight Agrobacterium culture was pelleted and resuspended into infiltration buffer (5% sucrose, 0.1% glucose, 0.1% MgSO4, 0.1‰ silwet, and 0.45 mM acetosyringone) to adjust O.D.600 to 0.4 by using. The resulting mixture was kept on ice for an hour before infiltration into tobacco leaves. After two days, the fluorescence signals were observed using Apotome.2 (Zeiss).

### Co-immunoprecipitation

The full-length coding sequence (CDS) of *HSBP* or *EID1* was amplified from cDNA of Col-0 seedlings and cloned into the entry vector, *pENTR/D/TOPO* (Invitrogen). The LR clonase II enzyme mix (Invitrogen) was used to transfer the insert to the destination vector *pEarleyGate 104* and *pB7-HFN* (Huang et al., 2016). The *pB7-HFN-EID1* and *pEarleyGate 103-HSBP* were transformed into the Agrobacterium GV3101 and transiently co-expressed in tobacco leaves using Agrobacterium infiltration. After 2 days, tobacco leaves were harvested and ground in liquid nitrogen, and the protein was extracted in Tris-protein extraction buffer (50 mM Tris-HCl, pH 7.4, 150 mM NaCl, 10% glycerol, 1% NP-40, 1 mM PMSF, 1× protease inhibitor cocktail (Roche), 5 μM EDTA, pH 8.0). The protein extracts were incubated at 4°C overnight with GFP-Trap beads (Chromo Tek, gtma). The immunoprecipitated samples were heated at 95 °C for 5min in SDS sample buffer (25 mM Tris-HCl pH 6.8, 2% SDS, 5% glycerol, 12.5 mM EDTA, 0.5% β-mercaptoethanol, 0.1 % (W/V) bromophenol blue). Protein separation was performed by SDS-PAGE, followed by immunodetection using mouse anti-FLAG (Sigma, F1804) and rabbit anti-GFP (Abcam, ab-290). The signal was detected with the iBright™ image system (Invitrogen).

### Generating transgenic lines

To generate the *EID1* complementation lines, the genomic fragment of *EID1* ( -1559 to +1008) was amplified with PCR using primers (**Supplemental Table S1**) and cloned into *pENTR-D-TOPO* vector (Invitrogen) and then subcloned into *pMDC204* vector (Curtis and Grossniklaus, 2003) to generate *pEID1::EID1-GFP* transgenic lines in the *eid1-2* background.

### The subcellular localization of HSBP with fluorescence confocal microscope

The full-length coding sequence (CDS) of *HSBP* or *EID1* were amplified from Col-0 cDNA and cloned into the entry vector, *pENTR/D/TOPO* (Invitrogen). The construct *pENTR-HSB*P served as template to generate the lysine mutations of *HSBP* using the Q5 Site Directed Mutagenesis Kit (NEB, #E0554S). The LR clonase enzyme mix (Invitrogen) was used to transfer the insert to the destination vector *pEarleyGate 104*. The primers were listed in **Supplemental Table S1.**

Constructed vectors were transiently expressed into Arabidopsis mesophyll protoplasts by polyethylene glycol (PEG)-Ca^2+^ mediated transfection (Yoo et al., 2007). The mCherry-NLS (pSAT6-mCherry-VirD2NLS, stock #CD3-1106) was used as a nuclear marker. YFP and mCherry fluorescences were observed using Leica TCS SP8 confocal microscope.

### GUS staining

For GUS staining assay, the *EID1* promoter was amplified the 1559 bp upstream of start codon and cloned into *pENTR/D-TOPO*, and transferred into *pMDC164* (stock #CD3-756) containing β-Glucuronidase reporter. The *proEID1::GUS* was transformed into Col-0 plants using Agrobacterium-mediated floral dip transformation method (Clough and Bent, 1998).

The 4-day-old *proEID1::GUS* transgenic plant seedings were treated without heat (CK) or at 37°C for 1 hour (H1) and recovered from HS for 1 or 2 hours (R1 or R2) individually. The assay was performed according to Guo et al. (2019) with modified. Briefly, the seedlings were soaked in GUS staining solution (0.1M NaPO4, 10mM EDTA, 0.1% Triton X-100, 0.5mM K_3_[Fe(CN)]_6_, 0.5mM K_4_[Fe(CN)]_6_, and 5mM X-Gluc) at 37°C for 3 hours, and wash the seedlings with 100% ethanol, then following with 70% ethanol. The Col-0 seedlings were used as a negative control for GUS staining. Staining was observed with a dissection microscope.

## Accession numbers

Sequence data from this article can be found in The Arabidopsis Information Resource under the following accession numbers: EID1 (AT4G02440) and HSBP (AT4G15802).

## Acknowledgments

We thank Dr. Tsung-Luo Jinn and Dr. Yee-Yung Charng for kindly sharing the *hsbp* and *hsp101* mutant seeds and for valuable discussion for the thermotolerance assays and manuscripts. We also thank the excellent technical assistance and the facilities of Technology Commons, College of Life Science, National Taiwan University.

## Author Contributions

The study was conceptualized by, Y.T.T., T.C.Y., G.L.C., and C.M.L.. The manuscript was written by C.M.L., Y.T.T., T.C.Y., and T.L. J., Y.T.T., T.C.Y., G.L.C., and Z.Q.W. designed, executed the experiments, and analyzed the data.

## List of supplemental data

Supplemental Table S1. The primers were used in this study.

Figure S1. The *eid1* mutants are knockout mutants.

Figure S2: The mRNA levels of endogenous *HSBP* in Col-0 and *eid1* mutants under heat stress.

## Funding

C.M.L.: NTU-CC-112L891804, NTU-CC-113L895003, NSTC-111-2311-B002-012-MY3

T.L.J.: NTU-CC-111L893002, NTU-CC-113L895002, NSTC-112-2311-B-002-020, NSTC-111-2311-B002-012-

T.C.Y.: PhD fellowship from NSTC, MOE, and NTU

Y.T.T.: NTU-CC-111L4000, NTU-CC-112L4000, NTU-CC-113L4000, NSTC-113-2811-B-002-122-

