## supplementary fig and table for "Arabidopsis EID1 E3 ubiquitin ligase regulates acquired thermotolerance by modulating HSBP translocation"

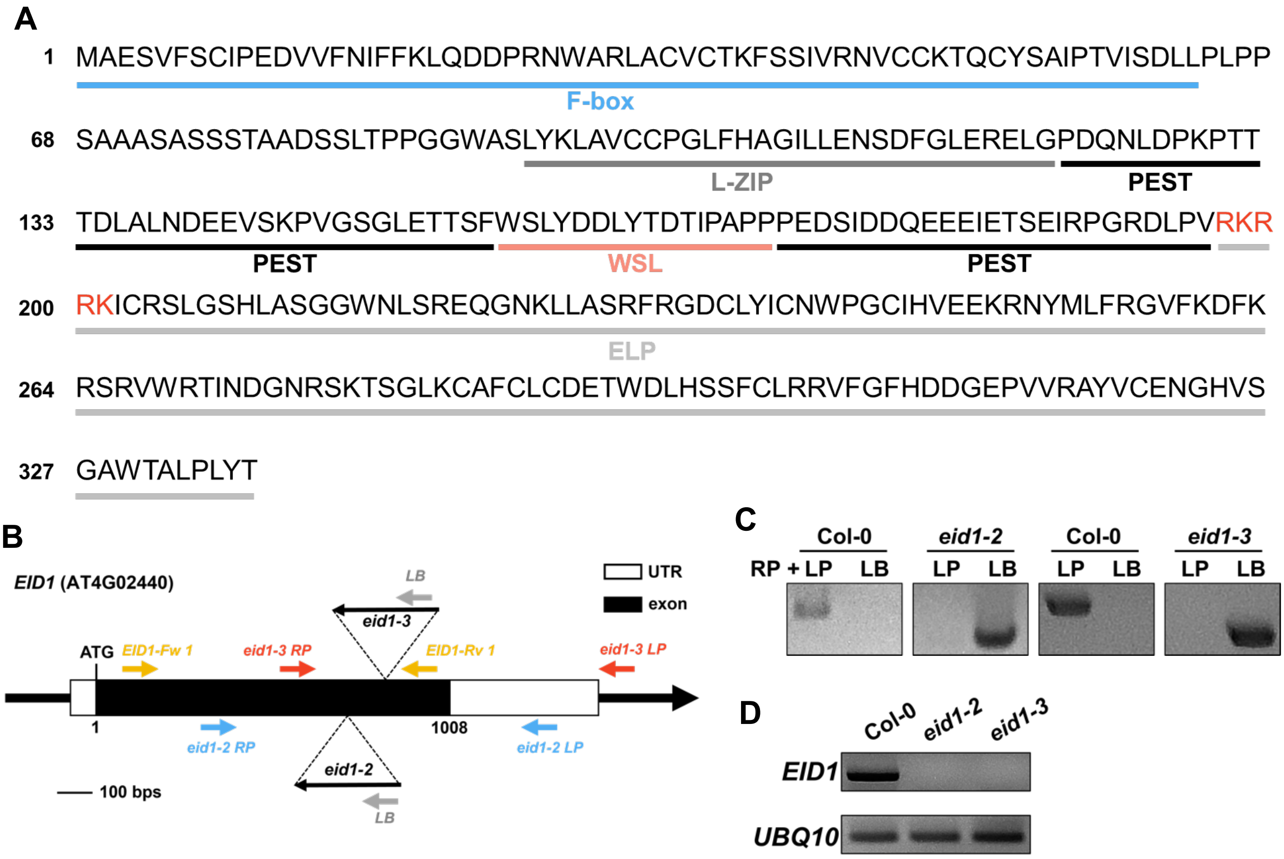


**Figure S1. The *eid1* mutants are knockout.**

**(A)** Scheme of EID1 protein sequence and its functional domains. **(B)** Scheme of T-DNA insertion site and primer sites of *eid1* mutants. The T-DNA insertions are in the exon (SALK_013061, *eid1-2*) and (SALK_027403, *eid1-3*), indicated as triangles, and their directions represented as arrows. The white and black boxes indicate UTR and exon, respectively. The yellow, red, blue, and gray arrows represent the primers used to confirm the T-DNA insertions and transcripts in *eid1* mutants. **(C)** Genotyping of *eid1* mutant alleles. The primer pairs of RP-LP and RP-LB were used to identify the wild type and T-DNA inserted alleles, respectively. **(D)** The transcript of *EID1* in *eid1-2* and *eid1-3* mutants by RT-PCR. *UBQ10* was used as an internal control.

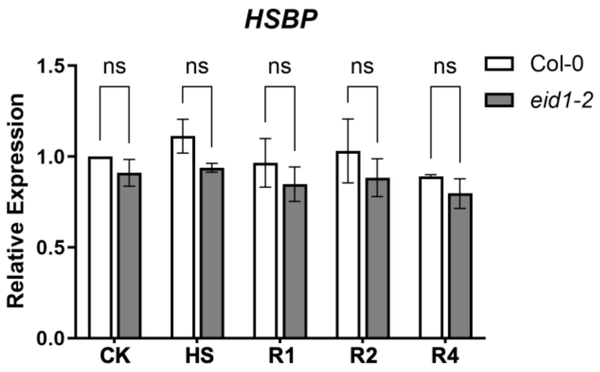


**Figure S2. The mRNA levels of endogenous *HSBP* in Col-0 and *eid1* mutants under heat stress.** The 4-d-old seedlings of *eid1-2* mutants without treatment (CK), heat-treat at 37 °C for 1 hr (HS), and following recovered 1, 2, or 4 hours (R1, R2, or R4) were harvested, and the *HSBP* transcripts were analyzed by real-time RT-PCR. The values are means ± SD of three biological repeats and asterisks indicate significant differences (ns, not significant, two-tailed *t*-test).

**Table S1: The primers were used in this study.**

| Name | Primer sequence (5' to 3') | Purpose |
|---|---|---|
| HSP17.6A-qF | TGTGAGGATGGAGAGGAGGATG | RT-qPCR |
| HSP17.6A-qR | CGTCATTACAAGCCGCAGAGAT | |
| HSP17.6II-qF | CAACGAGAAGACCCGCAACA | RT-qPCR |
| HSP17.6II-qR | GTCAGCAGGTGTAGCAGCCATT | |
| HSP17.6C-qF | CAAAACAGAGCAAACGCAAAGA | RT-qPCR |
| HSP17.6C-qR | AAACATCCAGCGAGAACGGA | |
| HSFA7a-F | GCATTCTTTCTCCACGATTCTCC | RT-qPCR |
| HSFA7a-R | GCAAATTCCCATCTCTCTGCTTC | |
| APX2 qFW | CTTGATGATCCTCTCTTTCTCCCA | RT-qPCR |
| APX2 qRv | ACTCCTTGTCAGCAAACCCGAG | |
| HSP18.2-qPCR-Fw | AAGGCAACAATGGAGAATGG | RT-qPCR |
| HSP18.2-qPCR-Rv | GCACACAAGCTTTTTATTTGACA | |
| HSP101-qPCR-Rv | ATGACCCGGTGTATGGTGCTAG | RT-qPCR |
| HSP101-qPCR-Fw | CGCCTGCATCTATGTAAACAGTG | |
| HSP70-qPCR-F | GGATGAGATATACAAAGGCGTGAA | RT-qPCR |
| HSP70-qPCR-R | AGGTGTGGCTTGTATGGTTAACAG | |
| EID1-qF3 | TTGTGCGATGAGACTTGGGATT | RT-qPCR |
| EID1-qR3 | TTGTGCGATGAGACTTGGGATT | |
| PP2AA3-qPCR-Rv | CCTGCGGTAATAACTGCATCT | RT-qPCR |
| PP2AA3-qPCR-Rv | CTTCACTTAGCTCCACCAAGCA | |
| pENTR-EID1-FW | caccATGGCGGAATCTGTCTTC | EID1 CDS cloning |
| pENTR-EID1-RV w/o STOP | AGTGTAGAGAGGTAAAGCAG | |
| pENTR-EID1pro-Fw | caccGTCACTGGGATGACTCCG | EID1 promoter cloning |
| pENTR-EID1pro-Rv | GGATCGCCTTCTTCTCTC | |
| pENTR-HSBP-Fw | caccATGGATGGTCATGATTCTGAGGATACTA | HSBP CDS cloning |
| HSBP-Rv | TTAAGAGGAACTAGCCGGTGTTTTGGGT | |
| pENTR-HSBP w/o STOP-Rv | AGAGGAACTAGCCGGTGTTTTG | |
| HSBP K10R-F | TGAGGATACTcgcCAGAGCACTGC | site-directed mutagenesis |
| HSBP K10R-R | GAATCATGACCATCCATTTAAG | |
| HSBP K41R-F | CATCATCACAcgcATTGATGACATGGGAG | site-directed mutagenesis |
| HSBP K41R-R | GAGTCCGACATTGTC | |
| HSBP K74R-F | TCCAGCCTCCcgcTCAGGCGATG | site-directed mutagenesis |
| HSBP K74R-R2 | GGAGGAGTGCCTTCTACTCCCATCTCGGCT | |
| HSBP K80R-F | CGATGAACCCcgcACACCGGCTAG | site-directed mutagenesis |
| HSBP K80R-R | AGGAGGAGTGCCTTC | |

| | | |
|---|---|---|
| SALK_013061_LP (*eid1-2* LP) | GCTTTTCAGGTTTATGAAACCC | *eid1-2* genotyping |
| SALK_013061_RP (*eid1-2* RP) | TGTTGTCCTGGTCTCTTCCAC | |
| SALK_027403_LP (*eid1-3* LP) | CCGTTAAGGAGGAAGGTCTTG | *eid1-3* genotyping |
| SALK_027403_RP (*eid1-3* RP) | TTTCGAAACCAGTTGGATCTG | |
| q-EID1-Fw2 (EID1- Fw-1) | CGATGAGACTTGGGATTTG | *EID1* in *eid1-2* and *eid1-3* |
| pENTR-EID1-RV w/o STOP (EID1- Rv-1) | AGTGTAGAGAGGTAAAGCAG | |
| UBQ10-Fw | TTCACTTGGTCTTGCGTCTG | RT-PCR |
| UBQ10-Rv | ATCTTGGCCTTCACGTTGTC | |
